# Blocking Osteoprotegerin Reprograms Cancer Associated Fibroblast to Promotes Immune Infiltration into the Tumor Microenvironment

**DOI:** 10.1101/2025.07.04.663190

**Authors:** Yao Wang, Hara Apostolopoulou, Im Hong Sun, Arjan Bains, David Gibbs, Sui Huang, Tamara Alliston, Ajay Maker, Thea Tlsty, Vasilis Ntranos, James M. Gardner, Anil Bhushan

## Abstract

The stromal compartment of many solid tumors plays a critical role in shaping an immunosuppressive microenvironment that limits the effectiveness of immune-based therapies^1^. Among stromal constituents, cancer-associated fibroblasts (CAFs) have emerged as key regulators of antitumor immunity^2–5^. Here, we identify a distinct subset of CAFs in both murine and human stroma-rich cancers that secrete osteoprotegerin (OPG)— a soluble decoy receptor that neutralizes receptor activator of nuclear factor kappa-B ligand (RANKL) and TNF-related apoptosis-inducing ligand (TRAIL), both of which are involved in T cell function. In vitro, OPG directly impairs CD8⁺ T cell-mediated killing of target cells. In murine models of pancreatic and breast cancer, antibody-mediated blockade of OPG promotes robust immune infiltration into the tumor microenvironment, leading to significant tumor regression. Stromal profiling revealed that OPG blockade induces a shift in CAF cells—reducing immunosuppressive OPG⁺ fibroblasts while expanding interferon-responsive fibroblasts, thus recalibrating the tumor stroma toward a pro-immunogenic landscape. These findings uncover a previously unrecognized mechanism of stromal immune suppression and highlight OPG as a stromal immune checkpoint controlling CD8⁺ T cell infiltration. Targeting OPG may offer a novel therapeutic strategy to convert immunologically “cold” tumors into T cell-infiltrated, tumor microenvironment.

## Introduction

Cancer cells induce alterations in the tissue microenvironment in which they grow, and the relationship between cancer and its microenvironment evolves throughout the life history of a tumor^1,6,7^. There has been considerable interest in understanding the relationship between the stroma and tumor-infiltrating immune cells, and emerging evidence suggests that the stromal compartment creates an immunosuppressive microenvironment that can also affect responsiveness to immunotherapy^8–10^. More recently, one cell type in the tumor microenvironment (TME), cancer-associated fibroblasts (CAF), have been prominently investigated ^3,11,12^. CAFs are not only a direct source of growth factors, but they can also release these factors via proteolysis of the extracellular matrix remodeling (ECM) by secreting the necessary proteases supporting primary tumor growth ^13–15^. The landscape of CAFs has been extensively profiled using a variety of techniques, such as single-cell RNA-seq and flow cytometry, across several mouse tumor models and human malignancies to uncover functionally distinct populations of CAFs ^5,16,17^. This has led to the increased recognition that CAFs are heterogeneous cell populations within the TME, and distinct subtypes play divergent roles, from supporting tumor growth to tumor-restraining functions – highlighting the limitations of designing nonspecific therapies broadly targeting all CAFs ^18^. Cancer-associated fibroblasts (CAFs) in solid tumors are functionally heterogeneous and can be broadly classified into two major subtypes: inflammatory CAFs (iCAFs) and myofibroblastic CAFs (myCAFs). iCAFs are characterized by secretion of inflammatory cytokines and growth factors and are driven primarily by NF-κB signaling; they often reside distally from tumor cells and are thought to promote immunosuppression via recruitment of myeloid cells^3,20^. In contrast, myCAFs exhibit high expression of α-smooth muscle actin (αSMA) and ECM components, are induced by TGF-β signaling, and are typically located adjacent to tumor nests, where they contribute to tissue stiffness and immune exclusion^19–23^. However, it remains unclear whether CAFs interact directly with immune cells to directly modulate adaptive immune-suppressive responses, and how we can harness the molecular mechanisms mediating this process to treat cold tumors.

Here, we identify a subset of inflammatory cancer-associated fibroblasts (iCAFs) characterized by the expression and secretion of osteoprotegerin (OPG; Tnfrsf11b), a soluble decoy receptor for TRAIL and RANKL. We show that iCAF-derived OPG directly impairs CD8⁺ T cell cytotoxic function via interference with TRAIL signaling. Using both genetic and antibody-mediated approaches to block OPG in murine models of pancreatic and breast cancer, we demonstrate that inhibition of this stromal-derived immune suppressive factor markedly enhances T cell infiltration and promotes tumor regression. These findings reveal a previously unrecognized mechanism of fibroblast-mediated immune evasion and highlight OPG as a stromal immune checkpoint that may be therapeutically targeted to overcome immune exclusion in stroma-rich, treatment-resistant cancers.

## Results

### Identification of osteoprotegerin (OPG)-secreting iCAFs

To probe stromal-immune interactions we first investigated an orthotopic breast cancer model using the GFP-labelled tumor cell line EO771 in C57BL/6J mice, as this immunocompetent neoplastic model is well established for interrogating the role of stromal-immune interactions during tumor growth^24^. To characterize the stromal microenvironment, tumors isolated on day 24 after implantation were subjected to flow cytometry sorting (GFP^-^CD45^-^), and scRNA-seq of the sorted stromal cells was performed using the Chromium (10X genomics) platform (Fig 1A). We excluded any remnant immune, endothelial and tumor cells, while using *Pdgfr* expression to selectively identify mesenchymal derived stromal cells. These stromal cells were clustered based on their individual transcriptomes and visualized by UMAP projection (Fig 1B). Using known gene markers, two distinct clusters that shared characteristics of CAF markers for fibroblasts, collagen deposition, and ECM were identified (Fig 1B, C, S1A, B). One of these clusters showed high expression of myofibroblast protein α-smooth muscle actin (gene *Acta2*) and *Pdgfrβ,* suggesting a myofibroblastic origin and were assigned as myCAFs based on widely used nomenclature (Fig 1C, D). The other CAF cluster was marked by high expression of *Periostin* (*Postn*) and the absence of *Acta2* ^25,26^. We also observed that a large fraction of the cells within this cluster displayed expression of cytokines such as IL33, chemokines such as Cxcl5, Cxcl12 and extracellular remodeling factors such as Mmp2 (Fig 1B, C, D). This combination of gene expression was reminiscent of a signature associated with activated fibroblasts or senescent cells, and based on previous nomenclature referred this Postn^+^ *Acta2*^-^ cluster as iCAFs ^8,27,28,29^. Of note, the Postn^+^ cluster is not homogenous, and only subset of cells within this cluster are post-mitotic (Fig S1C, D) ^30^. One key cytokine highly and uniquely expressed in this iCAF population is *Tnfrsf11b*, a member of the Tumor Necrosis Factor Receptor Superfamily (Fig 1D). *Tnfrsf11b encodes for a* secreted protein osteoprotegerin (OPG) that binds to and inhibits both receptor activator of nuclear factor kappa-B ligand (RANKL) and TNF-related apoptosis-inducing ligand (TRAIL) in cytotoxic T cells, to inhibit TRAIL-induced apoptosis ^31,32^. To establish whether OPG protein is directly secreted from CAFs, we next sorted CAFs from EO771 tumors using the scheme depicted (Fig 1E) and cultured them to collect conditioned media (CM). We detected high levels of OPG protein in CM from CAFs but failed to detect OPG from CM of epithelial and endothelial cells (Fig 1F). We also detected high levels of OPG from CM in sorted CAFs from human tumor samples, suggesting that both mouse and human tumors contain populations of iCAFs that secrete OPG (Fig 1G).

**Figure 1.**
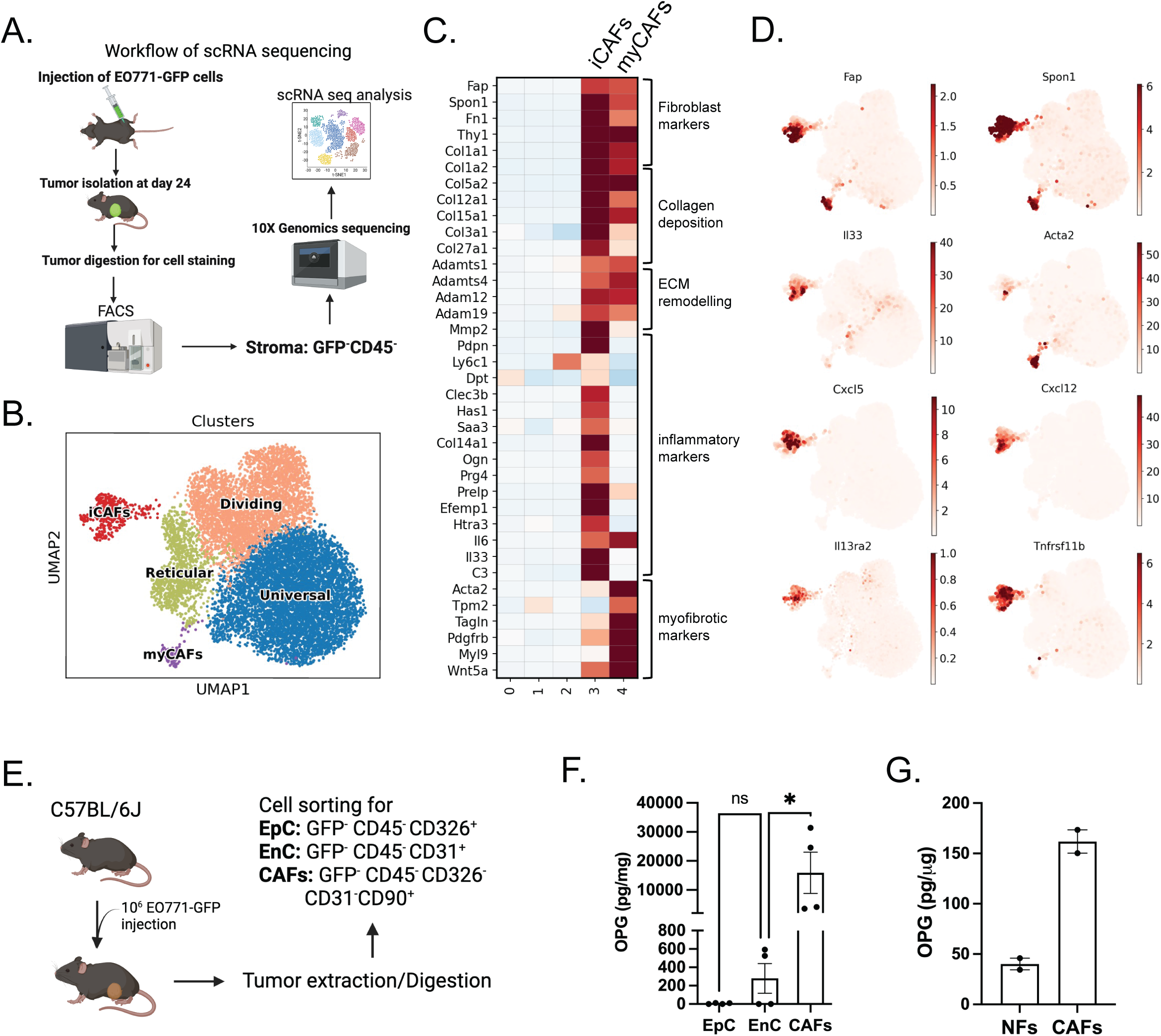
iCAFs secrete osteoprotegerin (OPG). (A-D) Data are from female C57/BL6 mice injected with GFP-labelled EO771 cells. Tumors were collected 24 days after implantation. Stromal cells (GFP^-^CD45^-^) were sorted by FACS and prepared for 10X Genomics scRNA sequencing analysis. (A) Schematic diagram of the experiment and single-cell RNA sequencing. (B) Uniform manifold approximation and projection (UMAP) visualization of immune-depleted stromal cells from tumor-injected mice generated from 10X Genomics single-cell RNA sequencing (scRNA-seq) analysis. Distinct clusters of stromal cells are identified using the Leiden algorithm and are represented by different colors. UF: Universal fibroblasts, DC: Dividing cells, RF: Reticular fibroblasts, *Postn+* CAFs: Cancer associated fibroblasts (CAFs) expressing high *Postn*, Acta2+ CAFs: CAFs expressing high *Acta2.*(C) Matrixplot showing high expression levels of literature-based CAF, iCAF, and myCAF biomarkers in clusters identified above as *Postn+* and *Acta2+* CAFs. Expression data were normalized with z-score transformation. Blue and red represent the low and high expression of a gene, respectively, relative to the median expression level. (D) UMAP plot highlighting the association between *Postn+* CAFs and an elevated “SenMayo score,” calculated based on the average expression of a curated set of 118 genes known to identify senescent cells at the single-cell level. (E). Schematic diagram of the cell sorting experiment (F) Quantification of OPG level from the sorted cells in F (G) Quantification of OPG secreted from nontumor-derived fibroblast (NFs) and cancer associated fibroblast (CAFs). NFs and CAFs were isolated from one patient with a gastric tumor by FACS using CD45^-^CD90^+^CD326^-^CD31^-^. Data in G are representative of two independent experiments. Statistics were calculated using ordinary one-way analysis of variance test.

To determine whether iCAFs in human stromagenic cancers also expressed TNFRSF11B, we analyzed stromal cells from human esophageal carcinoma using single-cell transcriptome sequencing from tumor samples and matched adjacent nonmalignant tissues from 12 patients ^33^. Multiple samples from each patient represented a path of progression from squamous epithelium through metaplasia, dysplasia to adenocarcinoma; to characterize the fibroblast cell diversity within samples, we performed integrated analysis restricted on this compartment across nonmalignant and tumor samples and identified five subclusters of fibroblasts (Fig 2A, S2A-C). Similar to the murine tumor model, in human esophageal carcinoma we identified cells with high expression of TNFRSF11B within the POSTN-expressing CAF cluster that also displayed expression of iCAF-characteristic transcripts, such as cytokines and chemokines (Fig 2A). We also analyzed stromal cell populations from a publicly available single-cell RNA-seq dataset of 26 treatment-naïve breast tumors (ER+, HER2+, and TNBC) from Wu et al. (2021) Fig 2 C)^62^. Using scVI for dimensionality reduction and Leiden clustering, we identified stromal subpopulations and visualized gene expression of *PDGFRA* and *TNFRSF11B* using UMAP projections (Fig 2 D). Overall, these results indicate that TNFRSF11B expression within CAFs is found in both human and rodent stromagenic cancers.

**Figure 2.**
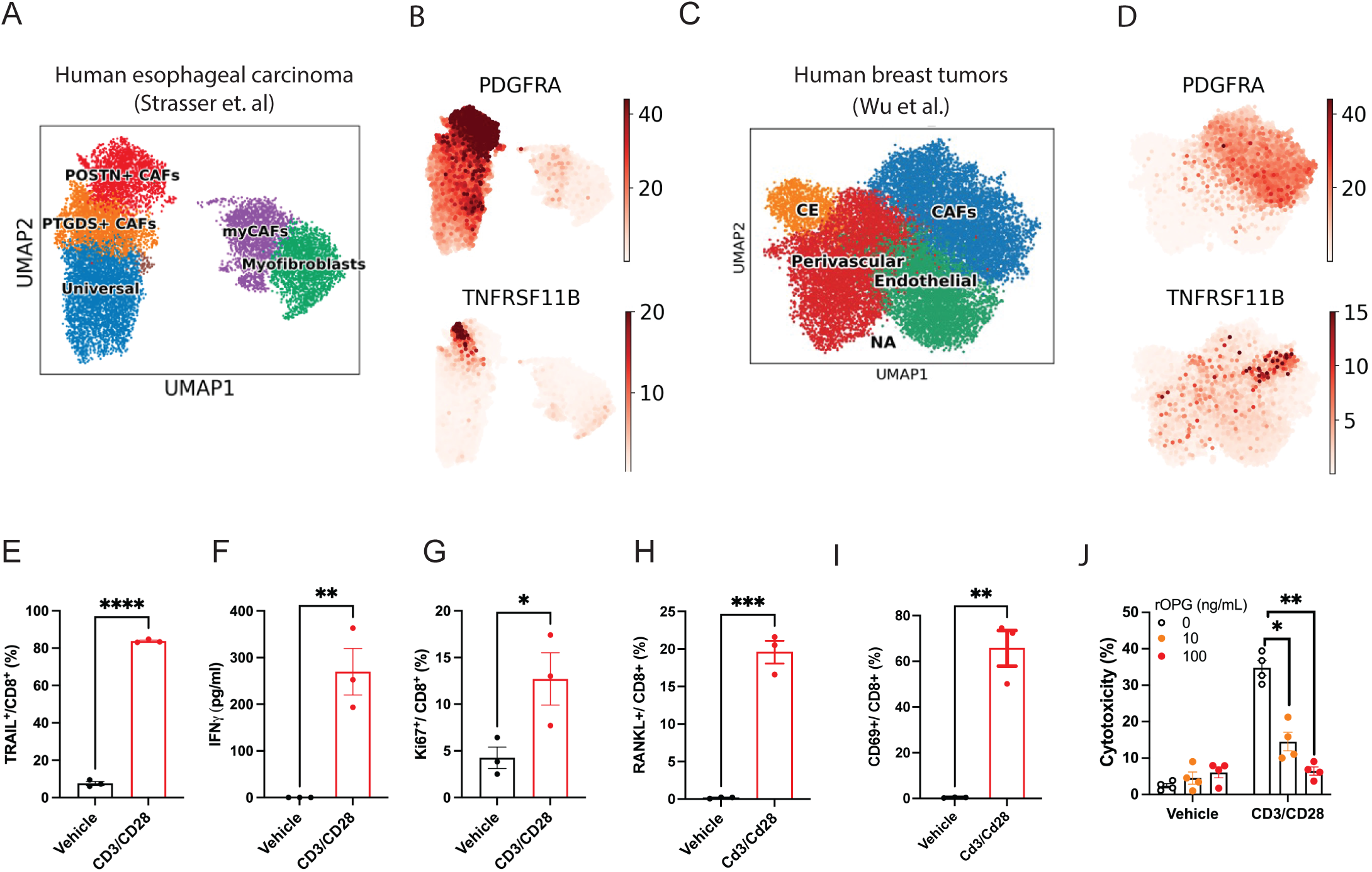
OPG inhibits cytotoxic function of T effector cells. (A–B) Data are from single-cell RNA sequencing (scRNA-seq) of 12 human esophageal carcinoma tumor samples, derived from biopsies or resections, as reported by Strasser et al., 2023. (A) UMAP visualization of different clusters of stromal populations. Distinct clusters were identified using the Leiden algorithm and are shown in different colors. Five fibroblast subtypes were identified: universal stromal cells, PTGDS⁺ cancer-associated fibroblasts (CAFs), myofibroblasts, POSTN⁺ CAFs, myofibrotic CAFs (myCAFs), and immune cells. (B) UMAP feature plots showing gene expression of PDGFRA (top) and TNFRSF11B (bottom). (C–D) Data are from scRNA-seq of 26 primary breast tumors, including 11 ER⁺, 5 HER2⁺, and 10 triple-negative breast cancer (TNBC) samples, as described by Wu et al., 2021. (C) UMAP plot showing four major clusters: CAFs, perivascular cells, endothelial cells, and CE (Cycling Endothelial). (D) UMAP feature plots displaying expression of PDGFRA (top) and TNFRSF11B (bottom). (E–L) Data are from isolated CD8⁺ T cells derived from mouse splenocytes, stimulated with vehicle or 3 μg/mL CD3/CD28 for 24 hours. (E) Quantification of TRAIL expression in CD8⁺ T cells. (F) Expression levels of IFNγ in CD8⁺ T cells, normalized to total protein. (G) Expression of Ki67 in CD8⁺ T cells. (H) Expression of RANKL in CD8⁺ T cells. (I) Expression of CD69 in CD8⁺ T cells. (J) Cytotoxic activity of CD8⁺ T cells against L929 cells at a target-to-effector (T:E) ratio of 1:4, with increasing concentrations of recombinant OPG (rOPG).

### OPG inhibits cytotoxic T-cell effector function

As OPG acts as a decoy protein that binds TRAIL and RANKL and inhibits its interaction with their corresponding receptors DR5 and RANK, we next explored whether OPG would directly interfere with immune mediated cytotoxic activity. Previous studies pointed to upregulation of TRAIL on the surface of T cells^61^. We first assessed whether CD8^+^T cells expressed TRAIL on the surface upon stimulation. Stimulation of effector CD8^+^ T cells, with CD3/CD28 antibodies led to a massive increase in cell-surface TRAIL expression (Fig 2 E); and the activation of CD8^+^ T cells was confirmed by increased IFN-γ secretion and proliferation (Fig 2 F, G). These results confirmed that TRAIL protein was upregulated on the cell surface of Teff cells upon activation. Similarly, stimulation of CD8^+^ T cells with CD3/CD28 antibodies led to an increase in cell-surface RANKL expression in a subset of cells that were activated as measured by CD69 upregulation (Fig 2 H-J).

To directly gain insight into whether OPG can block TRAIL-mediated cytotoxic activity of T cells we employed an effector-target co-culture assay using a TRAIL-sensitive fibroblast like cell line, L929 as the target cells, and sorted CD8^+^ T cells as effector cells from the mouse spleen. The cytotoxicity mediated by the Teff cells was measured by ATP levels as a proxy for target cell death. Co-culture of unstimulated CD8^+^ T cells with target cells did not result in cell death, however, when CD8^+^ T cells were stimulated with CD3/CD28 antibodies before co-culture, massive cytotoxicity activity was observed (Fig 2J). This cytotoxicity was dampened by the addition of recombinant OPG in a dose-dependent fashion (Fig 2J). To interrogate whether this same pathway operated in human cells we next assayed the ability of recombinant OPG protein to dampen the cytotoxic activity of human peripheral blood mononuclear cell (PBMCs)-derived CD8^+^ T cells and found a similar dose-dependent effect (Fig S2D-F).

### Blocking OPG alters CAFs

*As* αOPG affected CD8^+^ T cells function *in vitro,* we tested the effect of αOPG treatment *in vivo* in the orthotopic breast cancer model. GFP labelled-EO771 cells injected into the mammary gland of female C57BL/6J mice were treated with either αOPG or isotype control IgG (Fig 3A). Caliper measurements of the tumor size beginning two weeks after tumor injection showed that αOPG-treated mice displayed smaller tumors, and on average showed a 50% decrease in tumor size at the end of the four-week period (Fig 3B). The tumors dissected from αOPG-treated mice showed a 40% reduction in tumor volume on average compared to the IgG-treated mice (Fig 3C). To rule out off-target effects of αOPG and evaluate tumor growth kinetics and immune phenotype in the absence of OPG in the host, we next injected EO771 cells into female OPG-deficient and age-matched WT mice. Tumors in OPG-deficient (OPG KO) mice also grew more slowly and were reduced in volume compared to WT mice, recapitulating the phenotype seen in αOPG-treated mice (Fig 3D). To address whether the OPG secreting cells were derived from locally by tumor cells or derived from bone marrow, we carried our bone marrow chimera experiments in which bone marrow from OPG KO or WT age- and sex-matched littermates were transplanted into irradiated C57BL6-CD45.1(CD45.1) mice. Six weeks after radiation, half million of EO771 cells were injected into the 4th mammary fat of CD45.1 recipient mice and tumor growth were measured. No difference in the tumor size was observed in between recipient mice that received transplanted cells were from OPG KO or WT mice. This experiment indicated that OPG secreting stromal cells were derived from derived from local tissue resident stromal cells and not from bone marrow.

**Figure 3.**
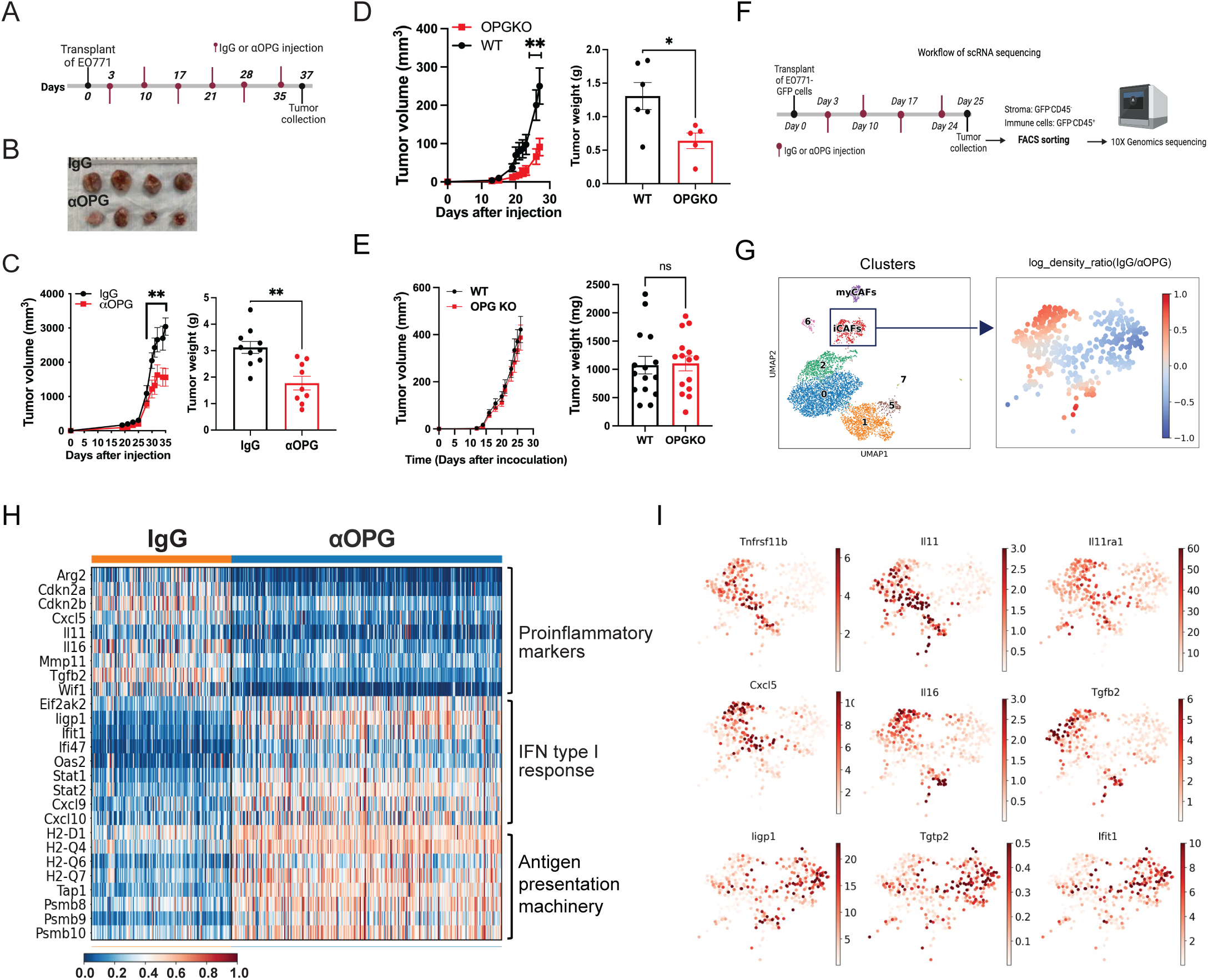
Blocking OPG alters tumor microenvironment. (A-C) Female p16^luc/+^ mice of 6 -8 weeks were used for EO771 injection. The mice were administered with 500 ng IgG or αOPG at 3, 10, 17, 28, and 35 days post-EO771 injection. (A) Schematic diagram of the experiment. (B) Tumor growth curve from IgG and αOPG-treated mice (n= 9 or 10 mice per group) (C) Tumor image (left)and weight (right) from IgG and αOPG-treated mice. (D) Tumor growth curve (left) and tumor weight (right) from age matched OPGKO and WT mice (N= 5 or 6 mice per group, The experiment has been repeated twice. (E) Irradiated C57BL6-CD45.1(CD45.1) recipient mice transplanted with bone marrow cells isolated from age- and sex-matched OPG KO or WT littermates. Tumor measurement was started at 12 days post tumor injection, and tumors were harvested at 26 days post tumor injection. (F-I) Female C57/BL6 mice were injected with GFP-labelled EO771 cells. Mice were treated with either αOPG or isotype control (IgG) at 3-, 10-, 17-, and 24-days post tumor implantation. Tumors were collected at 25 days post injection and processed for digestion and staining. Stromal cells (GFP^-^CD45^-^) and T cells (GFP^-^CD45^+^CD3^+^) were sorted by FACS and prepared for 10X Genomics scRNA sequencing analysis. (F) Schematic diagram of the experiment and single-cell RNA sequencing. (G) UMAP feature plots. UMAP plot of stromal cells from 10X Genomics scRNA-seq analysis (left). The stromal fraction was derived from mice injected with isotype control (IgG) or αOPG Colors indicate different clusters. UMAP plot showing high expression of *Postn* in cluster 3 (middle). Density plot showing the differences in density between IgG and αOPG treatment samples in Postn+ CAFs (cluster 3) (right). For each point in the UMAP space, calculated and visualized the likelihood of a UMAP region being associated with increased (positive log density ratio represented by red color) or decreased (negative log density ratio represented by blue color) proportions of IgG cells. (H) Heatmap showing relative expression levels of genes linked to inflamatory and Interferon (IFN) response pathway between IgG and αOPG treatment samples in *Postn+* CAFs. Expression data are scaled between 0 and 1, in which 0 (dark blue) and 1 (dark red) represent the minimum and maximum expression, respectively. (I) UMAP plots showing expression of key iCAF and interferon response pathway genes in *Postn+* CAFs. Data in B and C are representative of four independent experiments and D are representative of two independent experiments. Statistics were calculated using ordinary two-way analysis of variance test (B and D left panel) or two-tailed, unpaired Student’s t-test (C and D right panel).

Tumors from IgG- and αOPG-treated mice were dissociated and sorted into stromal (GFP^-^ CD45^-^) and immune (GFP^-^ CD45^+^) fractions and processed for scRNA-seq using the 10X platform (Fig 3F). The stromal cells were then subclustered and visualized by UMAP projection, enabling us to focus on the Postn^+^ *Acta2^-^*cluster that identified iCAFs (Fig 3F). A differential density map illustrates that cells originating from control IgG mice and αOPG-treated mice occupied distinct regions within the Postn^+^ cluster. Transcriptional profiling showed that expression of canonical markers of tumor fibroblasts, including *Pdgfra*, *Fap*, and *Pdpn* were indistinguishable between IgG treatment and αOPG-treated subsets (Fig S3C), but unlike iCAFs that originated from IgG-injected mice, iCAFs from αOPG-treated mice had an altered transcriptional gene profile (Fig 3F, G). Instead, pathway enrichment analysis pointed to the upregulation of IFN-responsive genes for MHC proteins and MHC-related molecules, suggesting that pathways regulated by IFN, such as antigen processing and presentation, were potentiated in the tumor fibroblasts that originated from αOPG-treated mice (Fig 3G, S3B). In addition, αOPG-treated fibroblasts upregulated type-I IFN-induced chemokines such as *Cxcl9* and *Cxcl10* that bind to the CXCR3 receptor expressed on CD4^+^ and CD8^+^ T cells (Fig S3C). Several studies have demonstrated a role for IFN-induced chemokines in the trafficking of T cells ^35–37;^ these results suggest that CAFs in the TME are modulated by αOPG treatment to enhance antigen presentation and activate chemokine secretion, with potential impacts on T-cell trafficking and activation within the tumor microenvironment.

### Blocking OPG enhances tumor infiltration and T cell effector function

The remodeling of the CAF landscape after αOPG-treatment, and in particular the appearance of a CAF subset with potential T-cell modulatory activity, prompted us to investigate the effect of αOPG-treatment on tumor T-cell infiltration (Fig 4 A). Spectral flow cytometry demonstrated the paucity of immune infiltration observed in isotype-treated tumors, in contrast to an extensive immune infiltration seen in tumors after αOPG treatment that included over a four-fold increase in CD4^+^ and CD8^+^ T cells (Fig 4B). The significant increase in immune infiltration in tumors after αOPG treatment was not a result of changes in tumor size (Fig 4SA), as these data were normalized to cell counts. We next examined the transcriptional changes in tumor-infiltrating lymphocytes (TILs) after αOPG treatment. UMAP density projection illustrated that CD4^+^ and CD8^+^ TILs that originate from control IgG treatment mice and αOPG-treated mice appeared to occupy distinct spaces representing distinct gene-expression programs (Fig 4C, S4B). Genes that control proximal signaling events downstream of the TCR, proliferation and T cell activation were upregulated in CD4^+^ and CD8^+^ TILs that originate from αOPG-treated mice when compared to isotype control-treated mice ^38^ (Fig 4C, S4C). Transcriptional analysis of TILs from αOPG-treated mice suggest features of enhanced effector functionality that could be harnessed.

**Figure 4.**
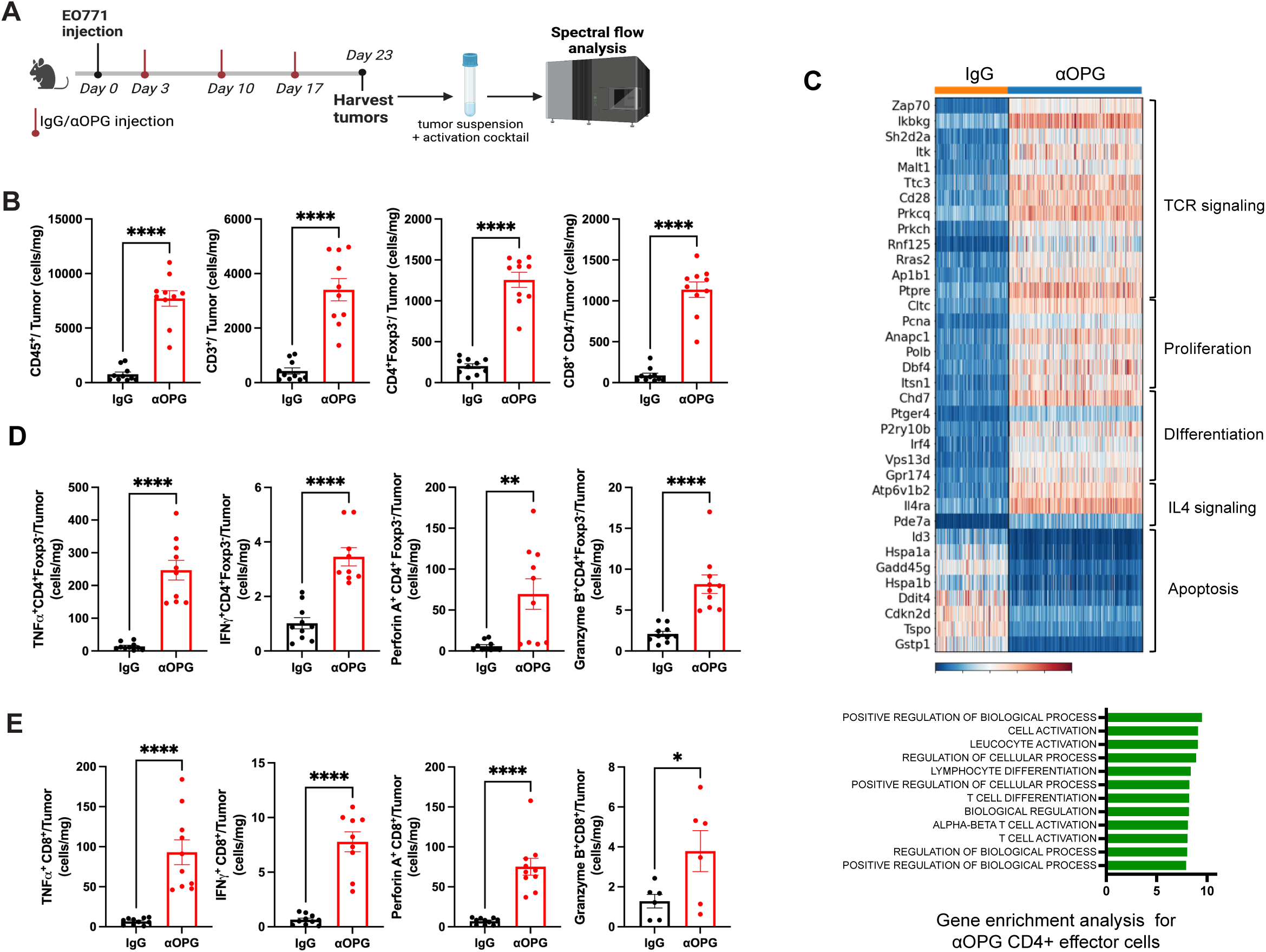
Blocking OPG enhances immune cell tumor infiltration and T cell effector function in breast cancer model. (A-E) Data are from C57/BL6 mice with 1×10^6^ EO771 implantation and treated with IgG or αOPG at 3-, 10-, and 17-days post implantation. Tumors were harvested 23 days post-implantation for T cell function assessment. (A) Schematic diagram of the experiment (B) Quantification of the total number of tumor infiltration immune cells (CD45^+^, CD3^+^ T, CD4^+^ effector T, and CD8^+^ T cells) normalized by tumor weight (n= 10 mice per group). (C) Heatmap showing upregulation of genes associated with T cell activation in CD4+ T effector lymphocyte upon αOPG treatment. Gene enrichment analysis show αOPG treatment resulted in T cell activation. (D) Quantification of the total number of TNFα, IFNγ, Perforin A, and Granzyme B on CD4^+^Foxp3^+^ T cells normalized by tumor weight (n =10 mice per group). (E) Quantification of the total number of TNFα, IFNγ, Perforin A, and Granzyme B on CD8^+^ T cells normalized by tumor weight (n =10 mice per group). Data in B, D and E are representative of four independent experiments. Statistics were calculated using two-tailed, unpaired Student’s t-test

However, extensive evidence has shown that transcriptional characteristics do not always directly translate into functional capacities; to test TIL effector functionality more directly, we next assessed the ability of TILs to secrete pro-inflammatory cytokines and release lytic granules (degranulation). We exposed single-cell suspensions of tumors from αOPG or isotype control mice to a cell-activation cocktail supplemented with Brefeldin A for four hours. Cells were subsequently stained for pro-inflammatory cytokines as well as perforin and granzymes indicative of cytotoxic function. Spectral flow analysis (Fig S4D) showed that stimulation resulted in dramatic increases in the staining of tumor necrosis factor alpha (TNF-α) and interferon gamma (IFN-γ) in CD4^+^ Foxp3^-^ and CD8^+^ T cells from αOPG-treated mice compared to isotype controls (Fig 4D, E). In addition to these pro-inflammatory cytokines, we also observed an increase in release of lytic granules containing perforin A and granzyme B from αOPG-treated samples, indicating enhanced CD4^+^ and CD8^+^ T cell effector function that could account for the tumor regression observed after treatment ^39^ (Fig 4 D, E). Spectral flow analysis of the tumors from *OPG-*deficient and WT mice also showed increased infiltration of TILs and was consistent with that from αOPG-treated mice compared to isotype controls (Fig S4E, F).

As OPG also plays important roles in bone remodeling, we additionally carried out trabecular bone analysis by microcomputed tomography (microCT) to evaluate bone loss over the course of αOPG and control IgG treatment^40^. After three weeks of treatment, no change was observed in trabecular bone volume/total bone volume (BV/TV) in αOPG treated mice compared to IgG-treated controls; we also measured bone-specific alkaline phosphatase (BALP), a serum biomarker for bone remodeling; no difference in BALP in the serum of αOPG and IgG treated mice was observed (Fig S4G, H). These data suggest that the three-week treatment of αOPG did not affect bone resorption or bone formation at detectable levels.

As CAFs that secrete OPG are present in several human stromagenic cancers, we next tested αOPG treatment in a pancreatic cancer model. FC1199 cells were implanted heterotopically into the flank of C57BL/6J mice (Fig 5A) and notably, the weights of tumors dissected from αOPG-treated mice showed a 30% reduction in tumor volume compared to IgG-treated controls (Fig 5B). F1199 tumor cells were dissociated, and immune infiltration and effector function of TILs analyzed by spectral flow cytometry. A significant increase in immune infiltration was observed in tumors after αOPG treatment in contrast to isotype-treated tumors, and moreover, TILs from αOPG-treated mice demonstrated an increase in release of lytic granules containing perforin A and granzyme B, indicating enhanced CD4^+^ and CD8^+^ T cell effector function similar to the results seen in the EO771 breast cancer model (Fig 5C).

**Figure 5.**
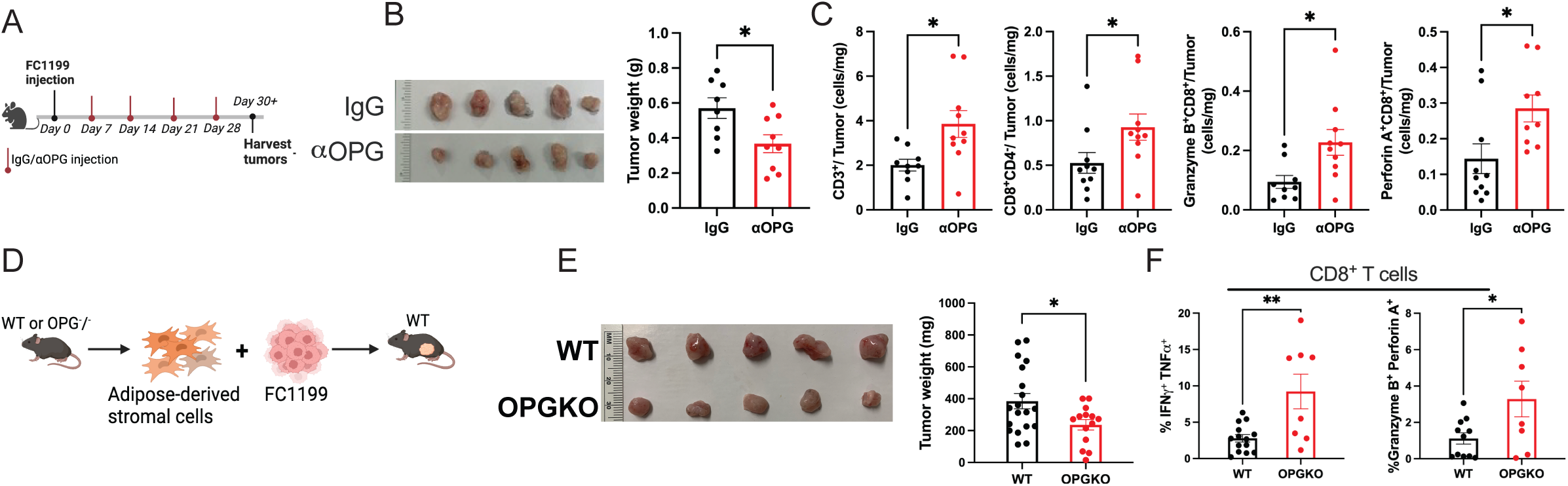
Blocking OPG enhances immune cell tumor infiltration and T cell effector function in pancreatic cancer model. (A-C) Data are from C57/BL6 mice injected with 1×10^6^ FC1199 and treated with IgG or αOPG at 7-, 14-, 21-, and 28-days post implantation. Tumors were harvested for T cell function assessment. (G) Schematic diagram of the experiment. (B) Tumor image (left) and tumor weight (right) on the day 34 post implantation (n=8 or 9 mice per group) (C) Quantification of the total number of CD3^+^ T cells, Cd4^-^CD8^+^ T cells, Granzyme B^+^, and Perforin A CD8^+^ T cells normalized by tumor weight (n= 8 or 9 mice per group) (D-F) Data are from C57/BL6 mice co-injected with 4×10^5^ FC1199 and 4×10^5^ adipose-derived stromal cells either from WT or OPG KO mice. Tumors were harvested at 25 days post co-implantation T. (D) Schematic diagram of the experiment. (E) Tumor weight on the day 25 days post implantation (n= 4 or 8 mice per group; combined two independent experiments). (F) Quantification of the frequency of IFNγ^+^ TNFα^+^ and Granzyme B^+^ Perforin A^+^ in CD8^+^ T cells (n= 4 or 8 mice per group; combined two independent experiments) Data in B, C, E and F are representative of two independent experiments. Statistics were calculated using two-tailed, unpaired Student’s t-test.

To determine whether OPG secreted specifically from iCAFs was the main source of immune evasion, adipose-derived stromal cells isolated from either WT or OPG KO mice combined with FC1199 were implanted heterotopically into the flank of C57BL/6J mice (Fig 5D). In these tumors, the microenvironment would predominantly contain CAFs derived from the co-implanted adipose-derived stromal cells. Analysis of these tumors revealed that those containing OPG-deficient stromal cells exhibited approximately a 35% reduction in weight compared to tumors containing wild-type (WT) stromal cells (Fig 5E). Consistent with the tumor reduction, CD8^+^ T cells displayed a significant increase in cytokines and lytic granules in the tumor with OPG deficient stroma in comparison to tumor that contained WT-derived stromal cells (Fig 5F). These results are consistent with the notion that blocking OPG secretion from CAFs primarily contributes to TIL activation and infiltration and also consistent with the bone marrow chimera experiments (3E).

## Discussion

It is increasingly recognized that immunosuppressive barriers within the tumor microenvironment are a major obstacle for checkpoint inhibitor and adoptive T-cell therapy in patients with solid tumors ^2,11^. This work identifies a cancer-associated fibroblast subtype in the tumor microenvironment that secrete an immunoregulatory protein, OPG, that acts as a critical negative regulator of T-cell tumor infiltration and effector function. Antibody mediated blockade of OPG or its genetic deletion in host animals induces massive infiltration of T cells that were capable of augmented cytokine secretion and effector function. These broad changes in the immune landscape are likely due to the alterations in the tumor microenvironment and the appearance of IFN-licensed CAFs, which play key roles in modulating immune infiltration to the tumor microenvironment ^12,41,42^. Moreover, immune infiltration and cytokine secretion can also lead to further activation of interferon-stimulated genes, creating a positive feedback loop that further enhances immune responses against the tumor. As immune infiltration of tumors in stromagenic cancers is closely related to clinical outcomes, a potential αOPG therapy that converts “cold” into “hot” tumors could improve treatment and survival, particularly when used in combination with other therapies ^43^.

While the evidence for a role of the CD8^+^ TILs in tumor control is compelling, our results indicate that CD4^+^ Foxp3^-^TILs also displayed the ability to secrete pro-inflammatory cytokines and release lytic granules in mice treated with αOPG. These results indicate that CD4^+^ Foxp3^-^TILs can exert cytotoxic functions and/or inhibit tumor growth through secretion of IFNγ and TNFa, consistent with previous work showing that CD4^+^ T cells play a substantial role in tumor control in both preclinical models and in patient case studies ^44^. While we have emphasized T cell infiltration into the tumor microenvironment, it is clear that infiltration of other immune cells such as myeloid cells are also observed in αOPG treated tumors.

Although interference with TRAIL/DR5 signaling plays a key role in restoring effector function in iNKT and T cells, many of the immunologic effects of αOPG are likely mediated by its modulation of the RANK/RANKL pathway. RANK/RANKL signaling appears to be involved in the emergence of IFN-licensed CAFs, and blockade of TRAIL alone has not replicated the robust immune activation observed with αOPG. In bone metabolism, OPG blockade enhances RANKL signaling and promotes osteoclastogenesis^40,45^. Given that RANKL signaling and T cells are known to influence cancer metastasis to bone, the mechanisms described here may have relevance to both primary and metastatic settings^46,47^. Interestingly, OPG blockade also leads to increased IFN-γ secretion by T effector cells, which may act in a negative feedback loop to regulate bone homeostasis—suggesting potential convergence between cancer immunology and bone biology, particularly in stromagenic cancers where bone loss is a common sequela.

It remains unclear whether the IFN-stimulated CAF subset that emerges after αOPG treatment arises through reprogramming of existing iCAFs, or from alternative lineage trajectories involving multipotent stromal progenitors. Our in vitro data indicate that TRAIL signaling can induce apoptosis of iCAFs, which would support a model in which IFN-licensed CAFs arise de novo from progenitor populations. This would further suggest that the primary targets of infiltrating T cells are iCAFs, rather than tumor cells. However, in vivo lineage tracing is needed to definitively resolve the origin and fate of these CAF populations.

Regardless of their ontogeny, our findings support a model in which OPG-directed therapies reshape the stromal-immune landscape of the TME, with broad implications for the treatment of otherwise refractory stromagenic malignancies.

## Materials and Methods

### Mice

Male or female C57BL6/J mice were obtained from The Jackson Laboratory (Cat# 00064). All mice were maintained under specific pathogen-free conditions, with free access to food and water, on a 12-h light-dark cycle. The sample sizes for each study are described in figure legends. All experiments were performed under the protocol approved by the Institutional Animal Care & Use Committee (IACUC) of University of California, San Francisco

### Cells

EO771, WI-38 cells, and L929 were obtained from ATCC (Cat# CRL-3461, CCL-75, and CCL-1) and cultured in DMEM supplemented with 1mM sodium pyruvate, 2mM HEPES, 10% fetal bovine serum (FBS), and 1% penicillin-streptomycin (Pen/Strep), or MEM supplemented with 10% FBS and 1% Pen/Strep, respectively. EO771 cells stably expressing GFP were generated using a GFP lentivirus. FC1199 cells were kindly gifted by Dr. Audrey Hendley and cultured in DMEM supplemented with 10% FBS and 1% Pen/Strep ^48^. Bone marrow-derived dendritic cells (BMDCs) were isolated from C57BL6/J mice and cultured according to the previously reported protocol ^49^. iNKT hybridoma DN32.D3 cells were kindly provided by Dr. Mitchell Kronenberg (La Jolla Institute for Allergy and Immunology) ^50^. iNKT hybridoma cells were maintained in the RPMI-1640 medium supplemented with 50 μM β-Mercaptoethanol and 10% FBS. Peripheral blood mononuclear cells (PBMCs) were isolated from Trima blood (Vitalant research institute) by using SepMate (STEMCELL technologies). All cell lines were tested for Mycoplasma contamination using MycoStrips from Invivogen. For all tumor injections, cells were used within passages 4-6.

### In vivo tumor models

For the orthotopic breast tumor model, EO771 cells were trypsinized, counted, and resuspended in PBS at a concentration of 2×10^7^ cells/mL. For C57BL6/J mice used, 8-10-week-old female mice were inoculated in the fourth mammary pad with 1×10^6^ cells in 50 μL. For *OPGKO* and wildtype (WT) mice, 8-10-week-old female mice were inoculated in the fourth mammary pad with 0.5×10^6^ cells in 50 μL. For the subcutaneous pancreatic ductal adenocarcinoma model, FC1199 cells were trypsinized, counted, and resuspended in PBS at a concentration of 1×10^7^ cells/mL. 8-week-old C57BL/6J male mice were subcutaneously inoculated in the right flank with 1×10^6^ cells in 100 μL. Flank skin hair was removed prior to implantation. Tumor volume (length and width) was measured using calipers with the modified ellipsoid formula: 0.5(length x width^2^). Tumor volumes larger than 4000mm^3^ were considered the endpoint, and those mice were removed. Mice with tumor ulcerations greater than 5mm were similarly removed from the study. When performing tumor measurements, the treatment conditions were blinded. For treatments, either OPG antibody (αOPG) or isotype (IgG) injection (i.p.) were injected on day 3, 10, 17, and 24 post cell injection for the EO771-injected mice, or on day 7, 14, 21, and 28 post injection for the FC1199-injected mice. In vivo-imaging data were analyzed using Living Image Software V 4.7.4. When harvested, tumors were weighed and processed for either single cell analysis by flow cytometer or SA-β-Gal staining analysis. All studies were approved by Institutional Animal Care and Use Committee of University of California, San Francisco.

### Bone marrow chimera

To generate OPG KO chimeric mice, C57BL6-CD45.1(CD45.1) recipient mice were irradiated with two 450 rads separated by 6 hr (Machine). Irradiated mice received 1 × 10^7^ bone marrow cells isolated from age- and sex-matched OPG KO or WT littermates. Six weeks after irradiation, half million of EO771 cells were injected into the 4th mammary fat of CD45.1 recipient mice. Tumor measurement was started at 12 days post tumor injection, and tumors were harvested at 26 days post tumor injection.

### Tumor isolation and digestion

Tumors were collected, weighed, and minced into 1mm^2^ pieces. Tumor tissue was digested with RPMI media containing 200 μg/mL collagenase P, 200 µg/mL DNAse I, and 100 µg/mL Dispase II, and digested as previously reported ^51^. After digestion, the cell suspension was passed through a 100 μm cell strainer, centrifuged at 400g for 10 min to obtain the pellet. Red blood cells were lysed by resuspending the pellet in 1 mL ACK lysis buffer and incubating for 3 min at RT. Lysis was attenuated by diluting the sample with complete cell culture media. The cells were counted using the Vi-CELL XR Cell Viability Analyzer (Beckman Coulter). Tumor single-cell suspensions were resuspended and processed for infiltration T cell function assessment.

### Cancer-associated Fibroblasts (CAFs) sorting and culture

Tumor single-cell suspensions were resuspended in flow cytometry buffer (1X PBS, 0.5% BSA, 2 mM EDTA), and blocked with Fc block (CD16/32 antibody, UCSF Antibody Core). After blocking, cells were stained with CD45-Pacific blue (Clone 30-F11), CD90-PE (Clone 53-2.1), CD31-PECy7 (Clone 390), and CD326-APC (Clone G8.8) for 30 mins on ice. Three different populations, epithelial cells (CD45^-^CD90^-^CD31^-^CD326^+^); endothelial cells (CD45^-^CD90^-^ CD31^+^CD326^-^); CAFs (CD45^-^CD90^+^CD31^-^CD326^-^) were sorted using BD FACS Aria II. Sorted cells were counted, adjusted cell concentration to 5 X 10^5^ cells/ mL and seeded into a 24-well plate. Supernatants were collected 24 hours post cell sorting and stored at -80 °C until use.

### OPG measurement

Mouse blood was harvested from the tail vein at the indicated time point. Samples were allowed to clot for 30 min at room temperature and then centrifuged for 20 min at 2,000g at 4 °C. Serum was collected and frozen at -80 °C until use. The supernatants of the cells were collected as indicated in the paper and stored at -80 °C for OPG measurement. The OPG levels were normalized to the RNA amount. OPG concentration was analyzed with an OPG ELISA kit. All samples were measured in duplicates.

### Splenocyte isolation

Mouse splenocytes were isolated using standard protocols. Briefly, mice were euthanized, and the spleens were excised and washed once in 1X PBS. After washing, the spleen was mashed through a 100 µm strainer with minimal force and the addition of 1X PBS. After that, the sample was strained through a 40 µm strainer and centrifuged at 500g for 5 min. The supernatant was discarded, red blood cells were lysed by resuspending the pellet in 1 mL ACK lysis buffer and incubating the sample for 4 min at RT. Lysis was attenuated by diluting the sample with 10 mL 1X PBS. The sample was centrifuged at 500g for 5 min, the supernatant was discarded, and the pellet was resuspended in 10 mL 1X PBS. Total live spleen cells were counted using the Vi-CELL XR Cell Viability Analyzer. Mouse CD8^+^ T cells were isolated from single-cell suspension of naive splenocytes by immunomagnetic negative selection using the MojoSort CD8^+^ T Cell Isolation Kit according to the manufacturer’s guidelines. CD8^+^ T cells were further seeded in triplicates, 8×10^4^ cells/well in 96-well plates, and treated with mouse CD3/CD28 dynabeads for 24 hours. After treatment, supernatants were collected for cytokine measurement by ELISA, and cells were stained and processed for flow cytometry.

### Adipocyte-derived stromal progenitor cells isolation from epididymal white adipose tissue (eWAT)

eWAT stromal cells were isolated using standard protocols. Briefly, mice were euthanized, and epididymal fat pads were harvested. The tissue was washed once in 1X PBS, minced with scissors, and digested with 1 U/mL Collagenase D in HBSS + 1% BSA for 30 min at 37 °C, with vigorous shaking every 10 min. Digestion was neutralized with 20 mL complete culture media (DMEM/F12+ 10% FBS + 1% Pen/Strep). The sample was strained once through a 100 µm cell strainer and centrifuged at 500g for 10 min at RT. The supernatant containing adipocytes and triglycerides was discarded from the top. Red blood cells were lysed by resuspending the pellet in 3 mL ACK lysis buffer and incubating the sample for 3 min at RT. Lysis was attenuated by diluting the sample with 20 mL complete culture media and centrifuged at 500g for 10 min at RT to obtain the eWAT stromal pellet. The supernatant was discarded, and the eWAT stromal cell pellet was resuspended in 1 mL complete culture media. Total live eWAT stromal cells were counted using the Vi-CELL XR Cell Viability Analyzer (Beckman Coulter).

### Co-injection of stromal cells and tumor cells

To investigate the significance of stromal-derived OPG in tumorigenesis, eWAT stromal cells were isolated from either age-matched OPGKO mice or WT mice, as mentioned earlier. The isolated eWAT stromal cells were washed with PBS and resuspended to a concentration of 4 × 10^6^ cells/ml. Simultaneously, the tumor cell line FC1199 cells were trypsinized and resuspended to a concentration of 4 × 10^6^ cells/ml. eWAT stromal cells and FC1199 cells were then combined at a 1:1 ratio immediately before injection. Subsequently, 10-week-old C57BL6 male mice were subcutaneously inoculated in the right flank with a 100 µL cell mixture, equivalent to 2 × 10^5^ eWAT stromal cells and FC1199 cells for each mouse.

### Cytotoxicity assay

The eWAT stromal cells was purified by CD45-depletion using mouse MojoSort mouse CD45 nanobeads. Senescence of eWAT stromal cells cells was induced by treatment with 20 μM etoposide for 24h. Proliferating eWAT stromal cells cells without etoposide treatment were used as the proliferative control (Pro). Bone marrow-derived dendritic cells (BMDC) were isolated from male C57BL6/J mice and cultivated. BMDCs were pulsed with 10 or 100 ng/L α-GalCer or vehicle overnight and then washed with PBS before co-cultivating with iNKT hybridoma cells at a 4:1 ratio. To test whether OPG impacts the cytotoxicity of T effector cells against target cells, eWAT stromal cells cells and L929 cells were pretreated with rOPG/αOPG and cultivated with iNKT hybridoma and CD8 T cells respectively for 8 hours. The viability of eWAT stromal cells cells or L929 cells was measured using ATPlite Luminescence Assay.

### Infiltration T cell function assessment

Tumor single cell suspension was adjusted to a cell concentration to 10^7^ cells/ml. 5×10^6^ cells were stimulated with cell activation cocktail (containing 0.081 µM phorbol-12-myristate 13-acetate, 1.34 µM ionomycin, and 5µg/ml Brefeldin A) at 37 °C for 4 hours. Post incubation, cells were washed twice with flow cytometry buffer, and once with PBS. Live cells were identified with Ghost-Red 780 Dye. Cells were blocked and stained using standard protocol as described previously. The following monoclonal antibodies were used for either cell surface or intracellular staining: CD3-APC (Clone 17A2), CD1d-Tetramer-PE (NIH Tetramer Core), TRAIL-PE/Cy7 (Clone N2B2), TRAIL-APC (Clone N2B2), NK1.1-BV711 (Clone PK136), FasL-APC (Clone MFL3), CD45-APC (Clone 30-F11), CD3-FITC (Clone 17A2), CD45-BV421 (Clone 30-F11), CD3-BUV737 (Clone 17A2), CD4-BV650 (Clone RM4.5), CD8a-BV605 (Clone 53-6.7), TNFα-PECy7 (Clone MP6-XT22), IFNγ-BV711 (Clone XMG1.2), Granzyme B-AF647 (Clone GB11), Perforin A-PE (Clone S16009A), and Foxp3-PECy5.5 (Clone FJK-16s). For intracellular staining, cells were fixed, permeabilized, and stained using the Invitrogen Fixation and Permeabilization kit according to manufacturer’s instructions. Data was acquired on the Spectral Flow Cytometer (Cytek) and analyzed using FlowJo. Data was processed using GraphPad Prism 9.

### Human peripheral blood mononuclear cells

Human peripheral blood mononuclear cells (PBMCs) were isolated from healthy patients using the SepMate PBMC isolation system (STEMCELL Technologies). The isolation of CD8+ cells was performed using the EasySep human CD8 negative selection kit (STEMCELL Technologies). Cell culture media consisted of RPMI-1640 containing 1% sodium pyruvate, 1% non-essential amino acid solution, 1% L-glutamine, 1% antibiotic antimycotic, 1% HEPES, and 10% fetal bovine serum (complete media). Wi38 cells 100 μM of etoposide was added to the media for 48 hours to induce senescence ^34^. Maintenance media was changed every other day for 6 days.

### Micro-computed tomography

Micro-computed tomography (μCT) was performed as previously described by CCMB core from UCSF^52^. Briefly, femur and tibia were isolated from either IgG or αOPG treated mice.

Measurement of volumetric bone density and bone volume at the right femur, tibio-fibular joint, or midshaft, and L5 vertebrae was using a Scanco Medical µCT 50 specimen scanner calibrated to a hydroxyapatite phantom.

### Serum bone alkaline phosphatase (BALP)

Mouse blood was harvested by cardiac puncture at the end of the experiment. Serum was isolated and stored at -80C before analyzed by ELISA kit.

### Statistical analysis

All data shown in this study represent one of three independent experiments, and the results are presented as mean ± SEM. Statistical analyses were performed with the two-tailed Student’s t-test for two-group comparisons, one-way ANOVA with Dunnett’s post hoc test for three- or four-group comparisons, or two-ANOVA. Data analysis was performed with Prism GraphPad (La Jolla, CA, USA). The levels of significance were set at *p < 0.05, **p < 0.01, ***p < 0.001, and ****p < 0.0001.

### Single cell RNA-seq processing and analysis

Single cell analysis of mouse stromal and immune cells from EO771-GFP tumors (Fig 1)

### Read processing and alignment

The sorted cells were assessed for viability using a hemacytometer and further processed for high-throughput droplet-based single-cell RNA sequencing (scRNA-Seq) using the 10x Chromium 3’ end platform (10X Genomics). Pseudoalignment with kallisto | bustools (Near-optimal probabilistic RNA-seq quantification^53,54^, Modular, efficient and constant-memory single-cell RNA-seq preprocessing.) was used to generate a gene count matrix. Initial preprocessing, including the removal of empty droplets and quality control using Scanpy (version 1.9.1) was performed ^55^.

### Quality control and normalization

Cells with less than 1000 detected counts and for which the total mitochondrial proportion exceeded 5% were excluded from further analysis. Doublets were then removed using scrublet ^56^ with default parameters. Genes with a minimum total count of 10 across all cells were also filtered out. The raw counts were normalized to 10,000 counts/cell and were log1p-transformed. For gene expression counts, the top 5000 highly variable genes were selected using the Scanpy function *sc.pp.highly_variable_genes*, employing Seurat flavor with default parameters ^55^. Subsequently, an scVI model was trained on the raw counts using all cells^57^. Clustering was performed using Leiden algorithm^58^ and visualization by using UMAP with default parameters^59^.

### Differential gene expression

Differential expression analysis across different clusters or between different treatment groups in the same cluster was performed using scVI^57^ All genes that were significantly upregulated (lfc_mean> 0, Bayes factor > 3) and expressed by at least 10% of the cells (non-zero proportions) were identified as markers.

### UMAP Differential Density Estimation (Fig 3)

We employed a Kernel density estimation approach to visualize the variations in cell subpopulation density between αOPG and IgG treatment samples. We separately calculated the densities of the αOPG and IgG samples using the UMAP coordinates of the respective cells, and then extrapolated the densities to the entire two-dimensional UMAP space using the nonparametric kernel_density.KDEMultivariate() function from the statsmodels Python package. At each point in the UMAP space, we calculated and displayed the logarithm of the estimated densities ratio, log (IgG_density/ αOPG_density), as a proxy for the likelihood of a UMAP region being associated with an increase (positive log density ratio) or decrease (negative log density ratio) in the proportion of IgG cells ^60^.

SenMayo score: Senescence was assessed using the scanpy.tl.score_genes function in Scanpy with a well-established gene set called SenMayo ^29^. SenMayo includes a collection of 117 genes in mice (Table S1) and 125 in humans (Table S2) that have been previously validated across various tissues demonstrating remarkable accuracy. We intentionally did not include TNFRSF11B to calculate “SenMayo score” as we wanted to confirm independently the correlation of TNFRSF11B expression and senescence.

### Filtering of cells

#### Stromal cells isolated from mouse EO771-GFP tumors (no treatment) (Fig 1)

The first scRNAseq dataset was obtained from stromal sorted cells isolated from an EO771-GFP tumor. To narrow down our analysis to fibroblast cells, we first excluded clusters with low expression levels of mesenchymal markers *Pdgfra* and *Pdgfrb*. Additionally, we eliminated one cluster containing low-quality cells, which exhibited significantly higher mitochondrial counts compared to the other clusters. As a result of these filtration steps, the final dataset consisted of 11,421 cells. We then reprocessed this dataset using the scVI method, visualized it using UMAP with 25 neighbors, and performed clustering with the Leiden algorithm using a coarse resolution parameter of 0.35.

#### Stromal and immune cells isolated from mouse EO771-GFP tumors treated with αOPG or IgG

This scRNAseq experiment included stromal (CD45-GFP-) and immune (CD45+GFP-) single cell suspensions derived from tumors dissected from mice injected with EO771-GFP cells and treated with αOPG neutralizing antibody or isotype control. Each library (αOPG/CD45-GFP-, IgG/CD45-GFP-, αOPG/CD45+GFP-, IgG/CD45-GFP-,) yielded high quality data, resulting in total 23072 cells. The initial data was split into two datasets based on Ptrpc expression: The immune dataset included Ptprc expressing cells and the stromal dataset included Ptprc non-expressing cells. The raw counts were reprocessed with scVI separately for the stromal and immune datasets using the same workflow as described above and visualized using UMAP with default parameters.

#### Single-cell analysis of human fibroblasts

Samples obtained from patients with Barret’s esophageal adenocarcinoma were processed, sequenced, and analyzed as in methods outlined in Strasser et al.^33^. Fibroblast-enriched scRNAseq data were acquired as raw counts and further processed. Genes with a minimum total count of 10 across all cells were filtered out. The raw counts were normalized to 10,000 counts/cell and then log1p-transformed. For gene expression counts, we identified the top 5000 highly variable genes using the Scanpy function sc.pp.highly_variable_genes, following the Seurat flavor with default parameters ^55^. Subsequently, we trained an scVI model on the raw counts using all cells ^57^. We used again the Leiden algorithm for clustering, resulting in a dataset with 5 clusters ^58^. We excluded cells with mtDNA ratio > 0.15 and one cluster characterized by low levels of mesenchymal markers PDGFRA and PDGFRB. We then reprocessed this dataset using the scVI method and visualized it using UMAP with default parameters ^59^. Clustering was conducted using the Leiden algorithm, and differential expression analysis across different clusters was performed using scVI ^57^.

---

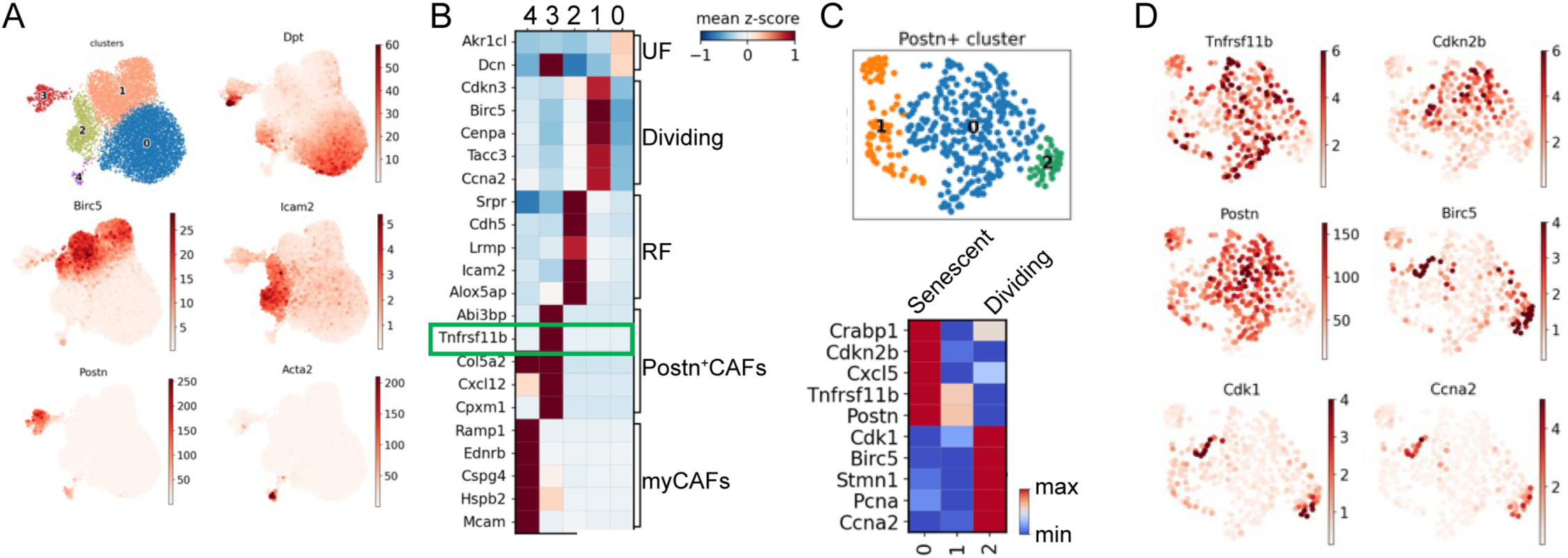

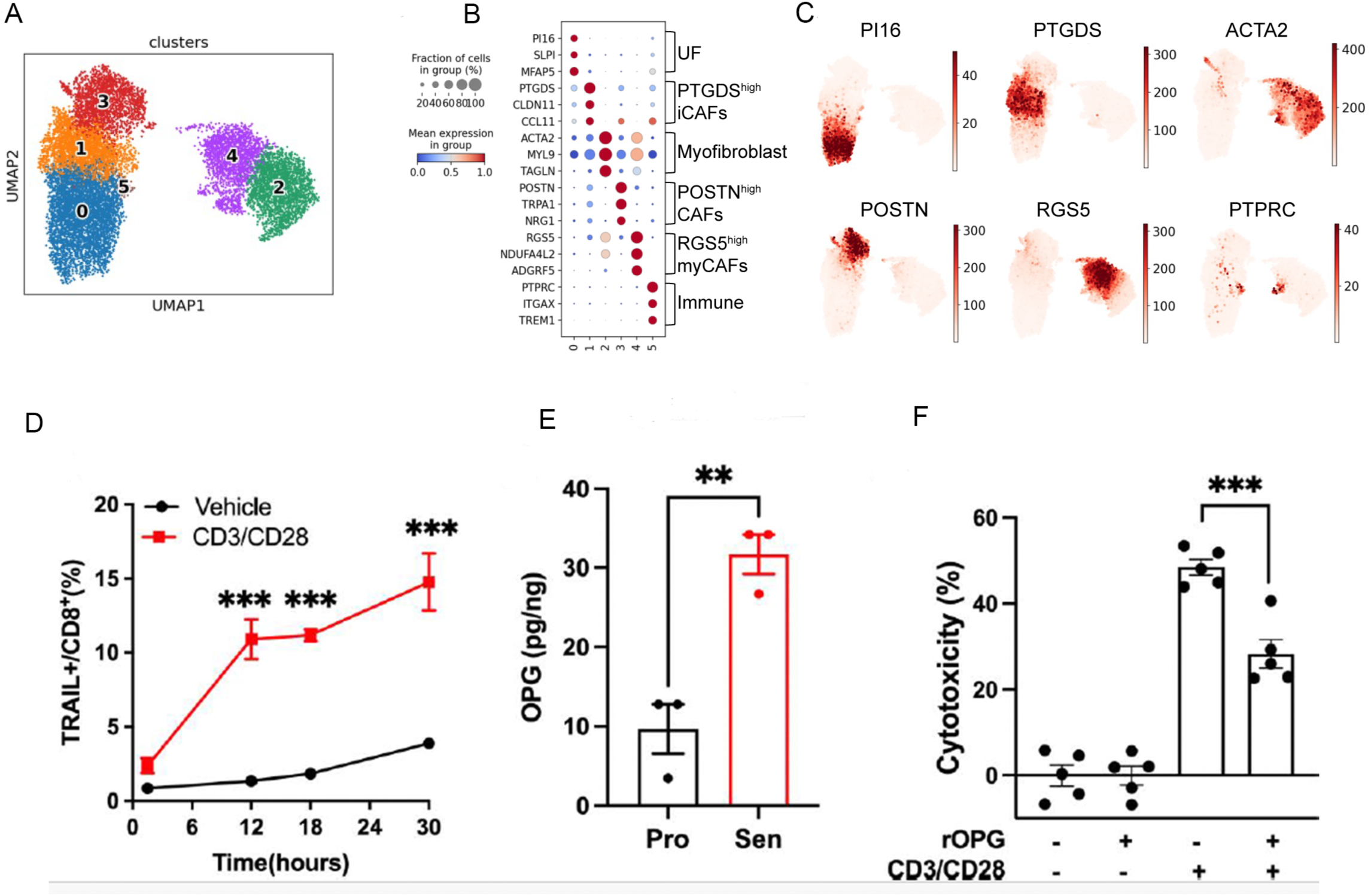

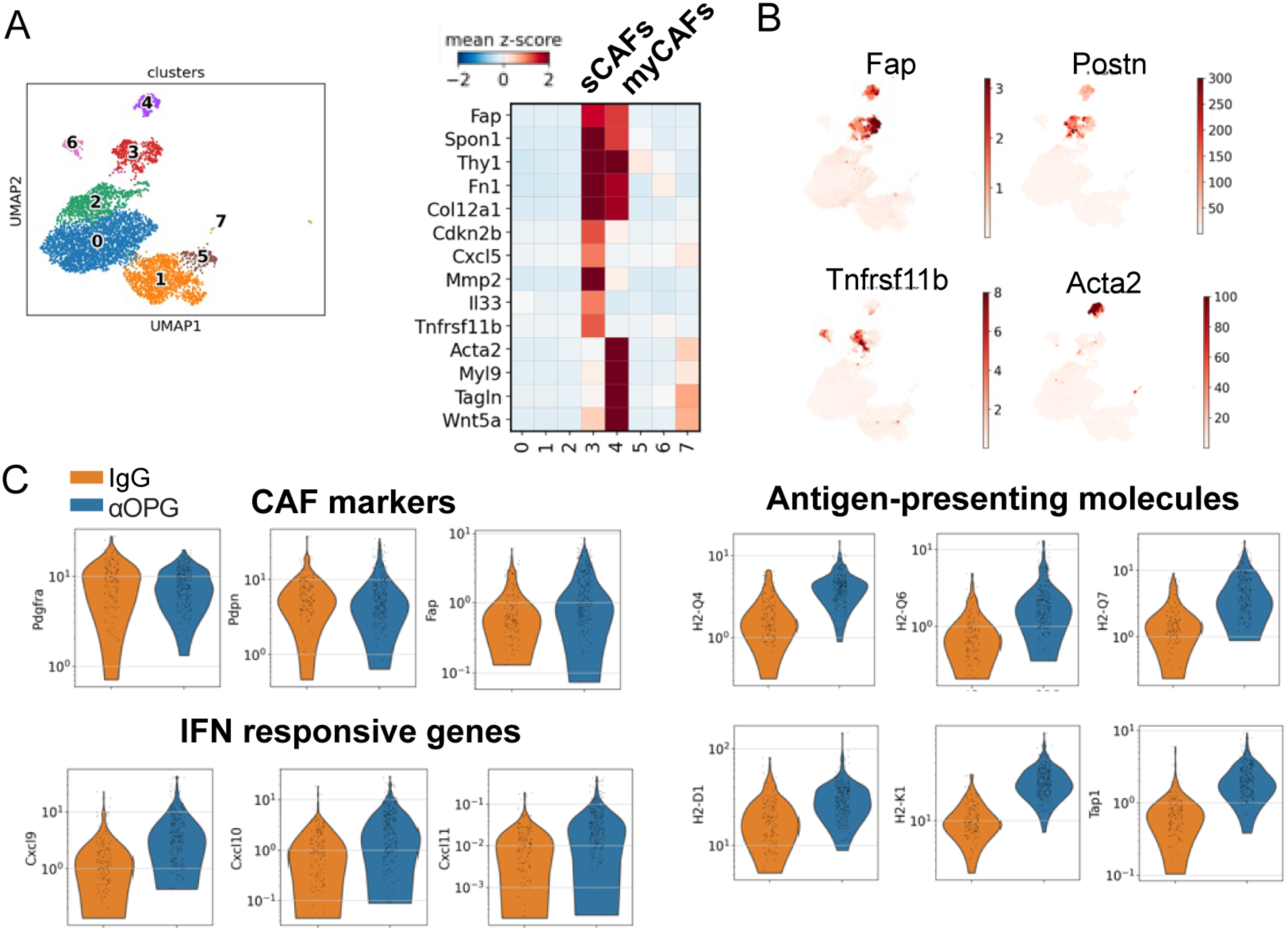

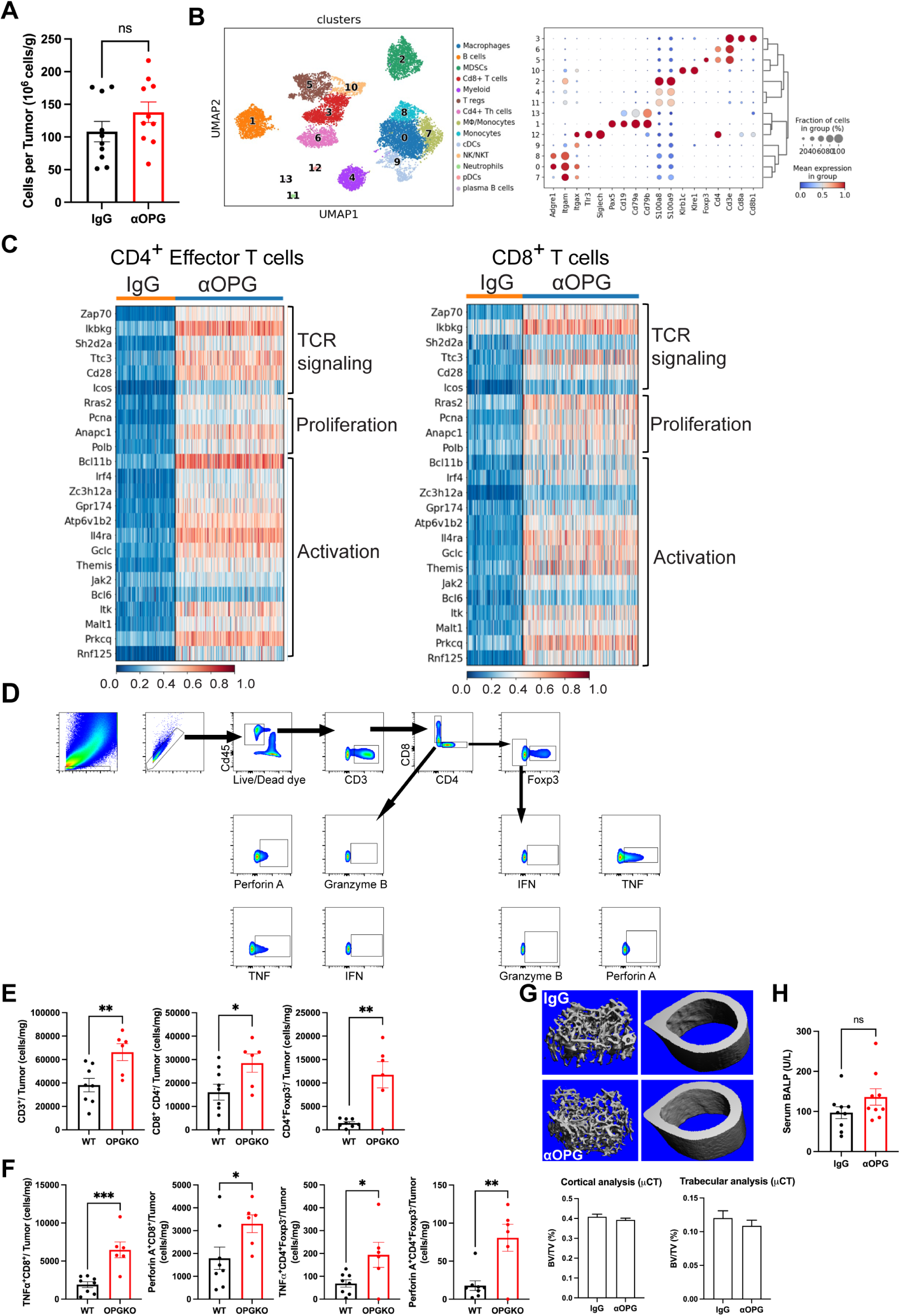

## Notes

### Competing Interest Statement

The authors have declared no competing interest.

